# Bistable State Switch Enables Ultrasensitive Feedback Control in Heterogeneous Microbial Populations

**DOI:** 10.1101/2020.11.10.377051

**Authors:** Xinying Ren, Christian Cuba Samaniego, Richard M. Murray, Elisa Franco

## Abstract

Molecular feedback control circuits can improve robustness of gene expression at the single cell-level. This achievement can be offset by requirements of rapid protein expression, that may induce cellular stress, known as burden, that reduces colony growth. To begin to address this challenge we take inspiration by ‘division-of-labor’ in heterogeneous cell populations: we propose to combine bistable switches and quorum sensing systems to coordinate gene expression at the population-level. We show that bistable switches in individual cells operating in parallel yield an ultrasensitive response, while cells maintain heterogeneous levels of gene expression to avoid burden across all cells. Within a feedback loop, these switches can achieve robust reference tracking and adaptation to disturbances at the population-level. We also demonstrate that molecular sequestration enables tunable hysteresis in individual switches, making it possible to obtain a wide range of stable population-level expressions.

## I. INTRODUCTION

Advances in synthetic biology have improved our ability to engineer genetic circuits and build microbes with controllable and complex, non-native functionalities. Processes such as biochemical synthesis of toxins and drug precursors often depend on population-level expression of one or multiple targets [1], [2], [3]. For optimal yield, these targets should be expressed while minimizing the influence of environmental fluctuations and disturbances [4]. Target expression should also be tunable, making it possible to tightly regulate a specific functionality, for example by balancing metabolic fluxes in biochemical production and drug dosage [5], [6].

Synthetic feedback circuits at the single cell-level have demonstrated robust and tunable gene expression in homogeneous populations [7], [8], [9], [10]. The inclusion of ultrasensitive modules in feedback circuits was recently shown to improve robust expression as a “high gain” feedback mechanism [11], [12]. A challenge posed by ultrasensitive mechanisms is that they may require transcription and translation of a large amount of components, imposing a major metabolic burden on the host cell, which may fail to thrive [13].

A possible route to mitigate the burden imposed by high-gain controllers at the single cell-level is the reliance on heterogeneous cellular states. Heterogeneity in gene expression levels is common in natural microbial populations, and often leads to diverse population phenotypes [14]. Importantly, heterogeneity in gene expression has been described as a strategy of ‘division-of-labor’ to relieve burden in single cells [15]. Further, heterogeneous populations better adapt to environmental disturbances, by taking advantage of sharp changes in phenotypical ratios that are induced by switching between distinct cellular states [16], [17].

In this paper, we outline a strategy study to use molecular controllers to control the concentration of a target protein in a heterogeneous microbial population. We show that an ultrasensitive transition in phenotype ratio can be achieved at the population-level by exploiting bistable switches operating in parallel in indivdual cells. We demonstrate that these bistable switches can be built using a positive feedback loop combined with molecular sequestration, which makes it possible to tune hysteresis. Finally, we illustrate how this heterogeneous population of bistable switches can be coordinated via quorum sensing, producing a population-level feedback system; collectively, the switches operate as an ultrasensitive controller that enables reference tracking and adaptation to disturbances at the population-level.

In the rest of the paper, we will use capital letters to indicate chemical species, and lower case letters to denote the the corresponding concentration. For example, species *X* has concentration *x*.

## II. ULTRASENSITIVE FEEDBACK CONTROL IMPROVES ROBUSTNESS IN POPULATION EXPRESSIONS

### A. Ultrasensitive Feedback Control

Ultrasensitive controllers can be used to achieve quasi-integral feedback in biological systems. Fig. 1A shows the steady-state diagram of an ultrasensitive controller and a process interconnected in a feedback loop [12]. The output *Y* of the process is the input to the controller, and the controller produces *U* as an output to actuate the process as an input. The steady state of the closed loop is determined by the intersection of input-output mappings of the controller and the process.

**Fig. 1.**
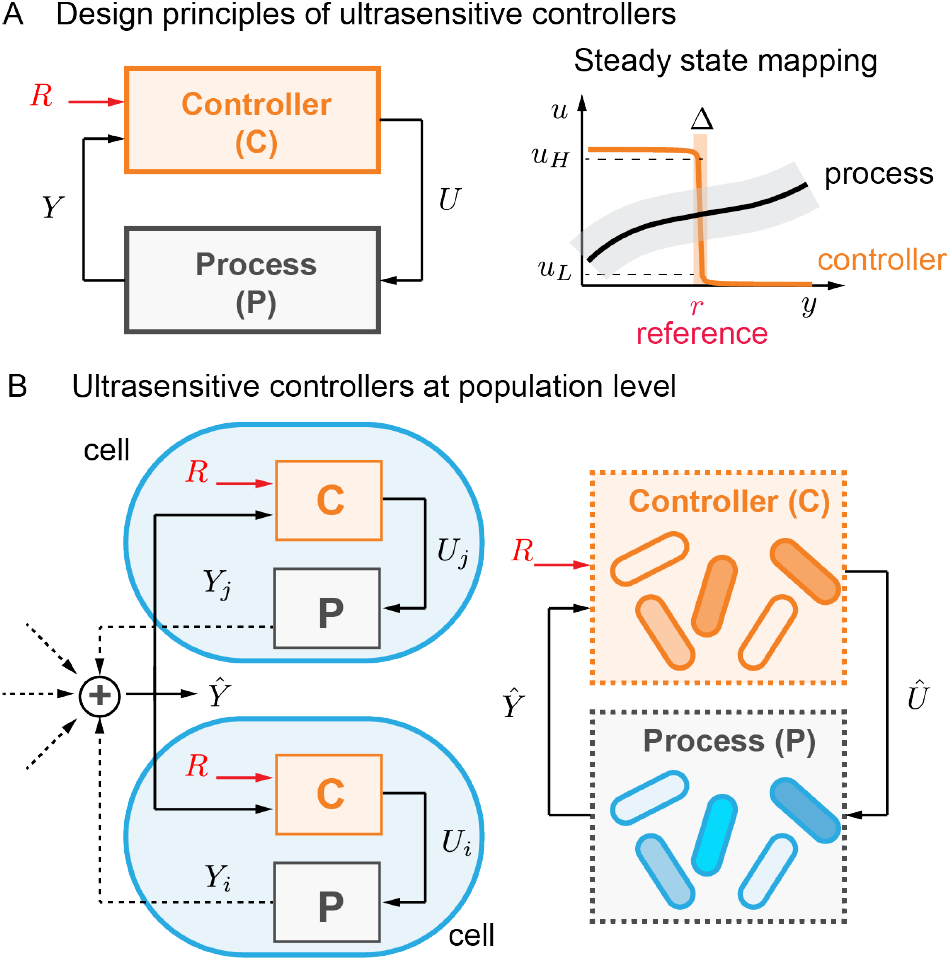
Ultrasensitive feedback controllers. Panel A presents a closed-loop diagram with an ultrasensitive feedback controller. The steady state input-output mapping demonstrates that the output is determined by the intersection of input-output maps and converges to the reference. Panel B is a schematic figure of a controlled system of a cell population. The process in each cell is regulated by a controller, and the population-level output is the total of individual cells. The population-level process can be considered under the regulation of a population-level controller with an overall *Û* and *Ŷ*.

When the controller is ultrasensitive, the input-output map of the controller exhibits a sharp transition, as demonstrated in Fig. 1A, right panel. Input-output maps of the controller (orange line) and the process (black line) intersect at the equilibrium of the closed-loop system. As long as the equilibrium is stable and lies in the ultrasensitve regime, the output *Y* defined by the intersection always converges to a neighborhood of the transition threshold, even when the process is uncertain or perturbed by disturbances (gray area). The ability to tune the threshold externally is analogous to setting the reference, and ultrasensitivity can be viewed as a “high-gain” mechanism that makes it possible to achieve quasi-integral behavior [12]. An important advantage of the concept of ultrasensitive controller is that it points to the individual roles of reference (threshold of the controller) and gain (slope of the controller) in a biological context. Further, it highlights that a multitude of ultrasensitive mechanisms could be used as quasi-integral feedback controllers [18].

Like other “high-gain” strategies, the implementation of ultrasensitive molecular controllers presents a potential challenge in that it may require large production rates of transcription and translation. Recent implementations of integral or quasi-integral controllers, relying for example on molecular sequestration or post-translational modification cycles, indicate that fast production of controller components may be necessary for correct operation [12], [19], [20]. A large production of proteins may induce cellular stress as it represents a burden on the transcription and translation machinery of the cell, and cells become stressed as a consequence. Therefore, it is important to explore strategies to take advantage of the concept of ultrasensitive feedback, without relying on drastic up-regulation of gene expression in individual cells.

### B. Ultrasensitive Controllers At The Population-Level

In many synthetic biology applications, the engineered functionalities of microbial cells are evaluated at the population-level. If synthetic circuits are directly designed at population-level, we might be able to avoid constraints at single cell-level, such as large production rates required for molecular controllers.

We consider a genetically-identical cell population of *n* cells. The control objective is the population-level expression of a target species *Ŷ*. Assume the i-th cell produces the target species at a concentration y_i_ . The population-level expression is considered as the sum of all single cells’ expressions, so we have the concentration of *Ŷ* as

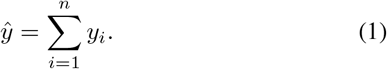

We assume all cells are under regulation of an identical type of synthetic feedback circuit, as shown in Fig. 1B, left panel. Each controller actuates the cell’s output *Y_i_* through input *U_i_*, and the controllers cooperatively drive the population-level output *Ŷ* to a reference *R*.

To perform population-level design of control circuits, we first define the population-level process and controller, as shown in Fig. 1B, right panel. Although physical implementations of circuits are based on biomolecular reactions inside each cell, the control mechanisms are better understood using a population-level description. The population-level process is defined as the sum of single cells’ processes, and is actuated by an overall control input *Û* and produces *Ŷ*. The population-level controller takes *Ŷ* as an input and compares it with the reference R and generates *Û* in output.

An ultrasensitive controller requires an ultrasensitive input-output response. We notice that a bistable switch circuit enables ON and OFF states with different gene expressions in a single cell, as shown in Fig. 2A. The hysteresis can generate a sharp switch in its input-output mapping. However, when combining a bistable switch with the process, the input-output maps’ intersection falls in the neighborhood of unstable equilibrium, leading to local instability in the closed loop, as shown in Fig. 2A, left panel. In addition, stochasticity of the cellular environment also causes a bistable switch to exhibit frequent state switching behaviors, as illustrated in Fig. 2A, right panel. Therefore, the bistable switch does not operate as a stabilizing ultrasensitive feedback controller in single cells.

On the other hand, if there are multiple bistable switches in parallel, we propose that it is possible to stabilize the total output by exploiting the sum of heterogeneous states of individual cells, as shown in Fig. 2B. The population operates as multiple single cells in parallel, thus bistable switches in individual cells can be considered as switches operating in parallel. Moreover, the stochastic state switching behavior in single cells does not interfere with the stable population-level expression, as shown in Fig. 2B, right panel. With this approach, bistable switch circuits can be used as an ultrasensitive controller at the population-level.

**Fig. 2.**
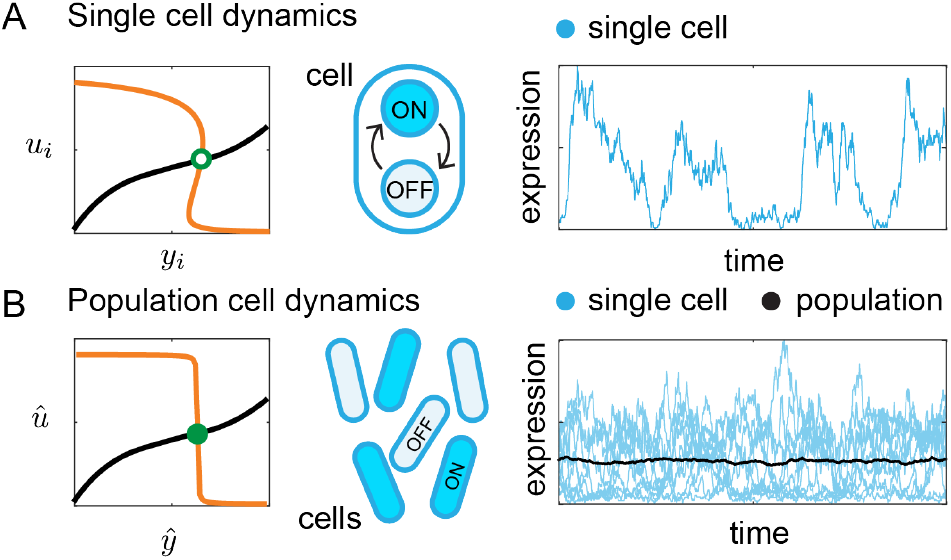
Ultrasensitive responses of bistable switches. Panel A shows the steady state input-output mapping of a single bistable switch and its dynamics. A bistable switch has two stable states (ON and OFF) and an unstable intermediate state. The intersection of input-output maps of the process and the single bistable switch results in an unstable equilibrium (empty green dot), and the stochastic trajectory of the single cell shows frequent state transitions. Panel B illustrates the population-level response and dynamics of multiple bistable switches tha operate in parallel. The steady state input-output mapping shows a stable equilibrium (filled green dot). The trajectory of population expression exhibits a constant output (black line).

In addition to the ultrasensitive response in the controller, it is necessary to design a mechanism to sense the total output, to coordinate switches in individual cells in a closed-loop system. We suggest that quorum sensing can be used to close the loop between the population-level process and controller. Quorum sensing molecules are secreted, diffuse, and mix in environments to form a global signal and activate downstream gene expressions in cells. Therefore, quorum sensing systems have been widely used in engineered microbial consortia to facilitate cell-cell communication and collaboration [21].

## III. Sequestration Generates Tunable Hysteresis In A Bistable Switch

First, we describe a bistable switch circuit with hysteresis in a single cell. A molecular bistable switch usually requires a positive feedback loop with high cooperativity. Recent studies have shown that sequestration with positive feedback is also sufficient to generate bistability [22]. In the following subsections, we consider a candidate bistable switch that combines positive feedback and molecular sequestration.

### A. A Single Bistable Switch With Molecular Sequestration

We present a circuit design including a self-activating species *X_i_*, sequestered by a species *Zi*, as shown in Fig. 3A. We assume the self-activation kinetics follows a Hill-type function, and production, degradation and sequestration follow the law of mass action. We can write down the model:

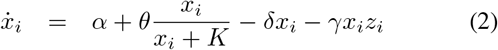

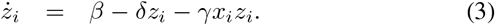

**Fig. 3.**
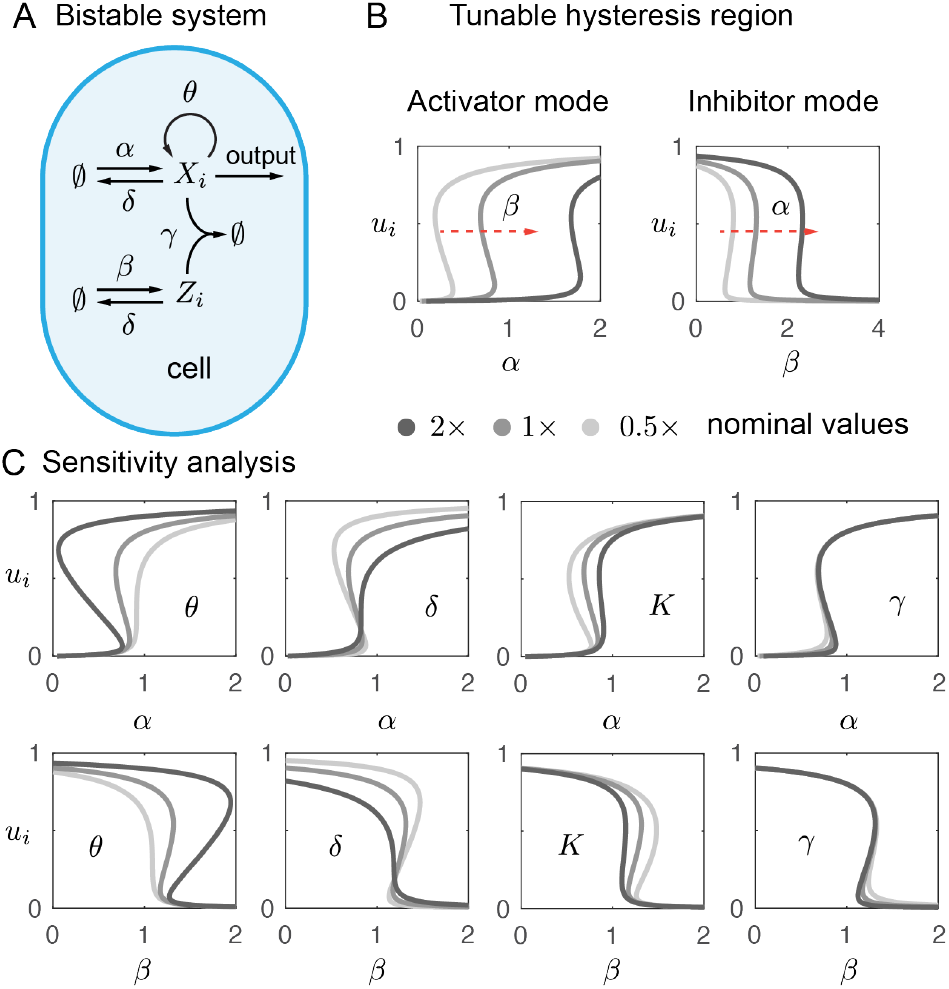
Tunable hysteresis enabled by molecular sequestration in a single bistable switch. Panel A illustrates the circuit design of a positive feedback loop coupled to a molecular sequestration mechanism. Panel B shows tunable input-output maps with varying *α* and *β*. Panel C demonstrates the effect of parameters on the input-output map. We use *K*= 0.2 for a better illustration in these sensitivity analysis plots.

The production rates *α* and *β* can be varied by external inputs, for example with inducible promoters.

We consider *α* and *β* as inputs to the bistable switch. When *α* is the input, we call it the activator mode, since the input activates *X_i_* production. On the other hand, the inhibitor mode corresponds to *β* being the input and inhibiting *X_i_* through sequestration. When the bistable switch is applied as the controller to the process, it should actuate the target expression. We assume such actuation is based on a Hill-type activation by *X_i_*, so we define the output of the bistable switch *U_i_*:

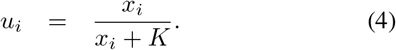

To test if the bistable switch has an ultrasensitive response, we derive closed-form solutions from a third order equilibrium equation that depends on parameters presented in Table I. We plot the steady state input-output maps in Fig. 3B for both activator mode (*u_i_* versus *α*) and inhibitor mode *u_i_* versus *β*). Both maps show hysteresis, indicating two stable equilibria (oN and oFF states) and one unstable equilibrium (intermediate state).

**TABLE I.** Simulation parameters.

| Parameter | Description | Value | Other studies |
| --- | --- | --- | --- |
| $\alpha, \theta, \beta, \rho$ ($\mu M \cdot \text{hr}^{-1}$) | Production | 1 | 0.1 – 100 [23], [24] |
| $\gamma$ ($\mu M^{-1} \text{hr}^{-1}$) | Sequestration | 500 | 36 – 3600 [25], [26] |
| $K$ ($\mu M$) | Dissociation | 0.1 | $10^{-5} - 1$ [27], [28] |
| $\delta, \phi$ ($\text{hr}^{-1}$) | Degradation | 1 | 0.36 – 3.6 [29] |

When the bistable switch is used as an ultrasensitive controller in the closed loop, the equilibrium should lie in the ultrasensitive regime, which is determined by the hysteresis threshold. Therefore, by tuning the threshold we can set the reference of the closed-loop output. To explore if the hysteresis is tunable, we change by two-fold the parameter values of *β* in the activator mode and *α* in the inhibitor mode, generating input-output maps in Fig. 3B. We find that both parameters move the hysteresis threshold in a broad range.

Next, we look for conditions on all parameters that ensure the bistability of this circuit analytically, and examine how parameters affect hysteresis properties with sensitivity analysis.

### B. Hysteresis Conditions And Sensitivity Analysis

First, to obtain hysteresis, there should be three distinct equilibria of equations (2)-(3). Since the variables represent concentrations of species, the solutions should be positive and real numbers.

We assume the sequestration between *X_i_* and *Z_i_* is fast [30], [31], i.e., 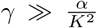. The analytical solutions can be approximated to find simple parameter conditions to admit three real roots (see Appendix A):

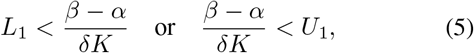

where

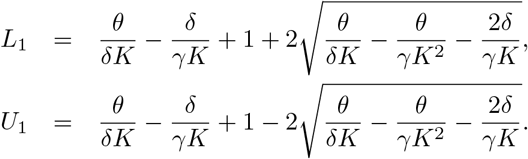

We use Descartes’s rule of sign to count the number of positive real solutions to admit three positive solutions (see Appendix A), which results in the following conditions:

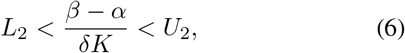

where 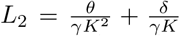, and 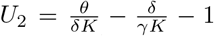. The conditions to admit three distinct roots that are real and positive are set by the intersection of conditions (32) and (33). Note that *U*_2_ < *L*_1_ then the conditions become

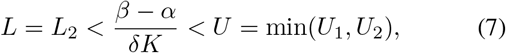

which can also be written as

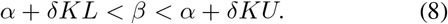

Equation (8) is a necessary condition for hysteresis. We first consider the inhibitor mode (*u_i_* versus *β*). Equation (8) determines the *β* regime that can generate three equilibrium solutions, which is the necessary bistability region. Given a fixed *α*, boundaries of the bistability region are set by *δKL* and *δKU*. We notice that L and U do not depend on α. It means that varying *α* only switches the the hysteresis threshold linearly without changing the left and right boundaries. This observation is consistent with input-output maps under different *α* values shown in Fig. 3B, right panel.

For the activator mode (*u_i_* versus *α*), we rewrite equation (8):

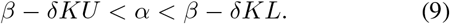

Similarly, the bistability region boundaries are set by *δKU* and *δKL*, and are not dependent on *β*. When varying *β*, only the hysteresis threshold is changed linearly, as shown in Fig. 3B, left panel.

Then we can inspect how parameters affect the bistability region by analyzing the sensitivity of boundaries *δKL* and *δKU* to parameters. For example, the sensitivity of *δKL* to parameter *θ* is defined as 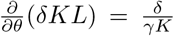. Similarly, we find sensitivity for other parameters: 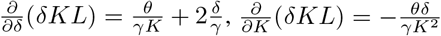 and 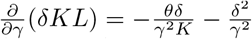.

We focus on the inhibitor mode (*u_i_* versus *β*) as an example of the analysis. We assume the sequestration rate is fast, where 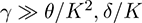, then *δKL* becomes insensitive to parameters *θ*, *δ*, *K*, *γ* since the sequestration rate *γ* is in the denominator of its sensitivity to all these parameters. It suggests that when we vary parameters *θ, δ, K* or *γ*, the left boundary of the bistability region will not present a large change, as also shown in Fig. 3C second row. On the other hand, the right boundary increases significantly with a larger *θ* and a smaller *δ* or *K*, according to the sensitivity analysis of *δKU* to these parameters. Similar conclusions can be drawn for the activator mode, which are consistent with input-output maps shown in Fig. 3C first row.

### C. Local Stability Conditions

Next, we study the local stability criteria of equilibrium to ensure there are two stable and one unstable solutions. We proceed to find the linearization of the system (2)-(3) and the Jacobian matrix *J* is derived as:

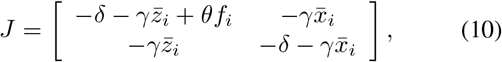

where 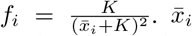 and 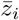 are the equilibrium. By assessing the sign of the real part of eigenvalues, we find the local stability is guaranteed when

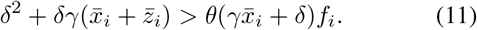

and local instability requires

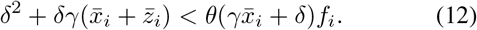

In summary, hysteresis conditions and local stability conditions determine the parameters for a bistable switch.

## IV. ULTRASENSITIVITY EMERGES FROM BISTABLE SWITCHES AT POPULATION-LEVEL

Now we evaluate the bistable switch circuit in a population-level setting, as shown in Fig. 4A. The population is considered as a collection of individual cells, where each cell has a bistable switch with the same input. They also generate an overall output *Û*. Therefore, bistable switches in parallel form the population-level controller.

**Fig. 4.**
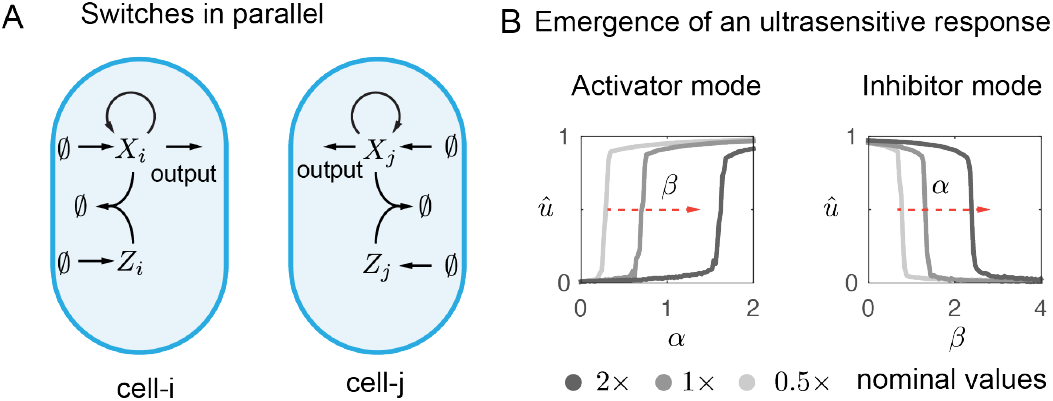
Multiple bistable switches in parallel. Panel A is a schematic figure of parallel bistable switches that represents a population of multiple cells. Panel B shows the population-level steady state input-output mapping of multiple switches. We simulate for *n* = 100 cells.

### A. Multiple Switches In Parallel

We expand the model to include *n* cells with bistable switch circuits. The dynamics of all cells are

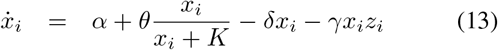

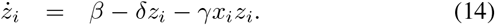

for *i* = 1: *n*. Since the population-level controller generates an actuation to the population-level dynamics through the overall activation in *Ŷ* production by *X_i_* in all cells, we define the total output of multiple bistable switches *Û*:

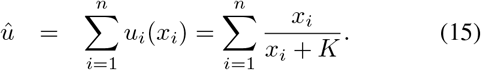

Again, we consider *α* and *β* are inputs to all bistable switches. Then we evaluate the steady state input-output maps: *û* versus *α* for the activator mode and U versus *β* for the inhibitor mode, by numerical simulations. As shown in Fig. 4B, the population-level input-output map exhibits a graded increase or decrease along with the varying parameters. In contrast to a single bistable switch, multiple switches in parallel reach stable equilibrium in the ultrasensitive regime. Meanwhile, the thresholds of the graded maps are tunable by changing parameters *β* and *α*.

### B. Graded Output At Population-Level

The operation of multiple switches in parallel results in emergent properties of the input-output map. As shown in Fig. 4B, while individual switches are bistable, the total output exhibits a graded response to the input, and all total outputs admit a stable level.

Recall that a single bistable switch only presents two stable states (oN and oFF states), and one unstable state (intermediate state). The output of multiple switches is the sum of all single switch’s states. Whenever a single bistable switch changes its state, the total output admits a new stable equilibrium. Since the numbers of single switches in oN states and oFF states are no longer restricted, the total output can reach a larger range of stable equilibrium. Therefore, in a population that consists of millions of cells, the extreme large number of bistable switches in parallel enables a smooth and graded response with stable outputs.

### C. Emergent Ultrasensitivity At Population-Level

Multiple switches also generate an ultrasensitive inputoutput map at population-level, via a sharp transition of the oN/oFF population ratio. The ultrasensitivity emerges from the sharp transition in single switches. To better understand how transition rates between oN and oFF states effect the ultrasensitive response, we consider a simple population-level model of multiple switches:

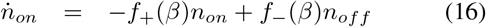

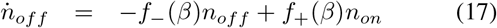

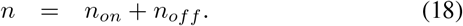

Variables *n_on_* and *n_off_* are total numbers of cells that exhibit ON and OFF states, and f_+_ and f_are transition rates from oN to oFF and vice versa. Here we focus on the inhibitor mode given a fixed α, so the transition rates depend on the input *β*. At steady state, we have

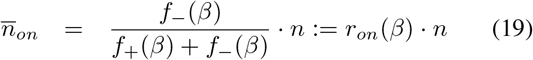

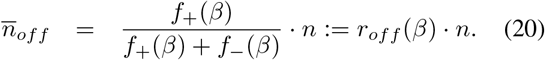

Variables 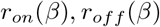 are defined as population ratios of ON and OFF cells at steady state.

For simplicity, we assume a bistable switch at ON state generates an output *U_on_*, and at OFF state it generates an output *U_off_*. As defined in equation (4), the output of a single bistable switch depends on *X_i_* concentration. Since the concentration of *X_i_* is determined by input *β* in the inhibitor mode, we use *U_on_*(*β*) and *U_off_* (*β*) to represent the output values of ON state and OFF state switches in cells. According to equation (15) and equations (19)-(20), we can derive the population-level output *Û* at steady state as

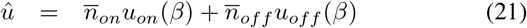

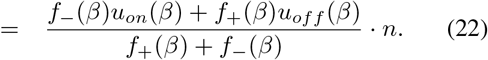

It shows that the population-level output *Û* of bistable switches not only depends on the single cell-level output, but also transition rates between states. We can find out in Fig. 3B, right panel, that the outputs at either ON state or OFF state are not very sensible to *β* and have rather flat curves compared to the intermediate transition. Meanwhile, transition rates 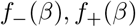 can be very sensitive to *β* in bistable switches. The population-level input-output response becomes ultrasensitive because of sensitive transition rates.

We can also rewrite equation (22) with population ratios *r_on_* and *r_off_*:

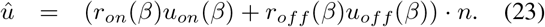

If we assume OFF state generates a very small output, i.e., 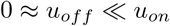, we can obtain

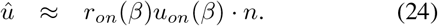

It implies n ultrasensitive controller can be achieved by sharp population ratio changes. In other words, if *r_on_*(*β*) is ultrasensitive to *β*, the output u becomes ultrasensitive, which only appears at population-level. Such emergent properties suggest that bistable switch circuits can be used for ultrasensitive control at population-level without requiring large productions in single cells.

## V. Quorum Sensing Coordinates Bistable Switches In The Closed Loop

Finally, we use bistable switches as a collective ultrasensitive feedback controller to control population-level expression. We use a quorum sensing system to link the population-level process and controller. Quorum sensing signals can coordinate cells’ state switching behaviors according to the error between the population-level output and the reference.

For simplicity, we consider the desired population-level output is the concentration of the quorum sensing signal. As shown in Fig. 5A, we adopt the inhibitor mode and link the output *Ŷ* to the activation of *Z_i_*, forming a negative feedback loop. A synthetic circuit implementation is also proposed in Fig. 5B. We suggest that a sigma factor activates itself and an enzyme LuxI that catalyzes the synthesis of a quorum sensing signaling molecule AHL. The signaling molecule diffuses across cell membranes and activates an anti-sigma factor that can sequester the sigma factor and form an inactive complex. There is another inducible production of the sigma factor, which can be used to set references by external inducers.

**Fig. 5.**
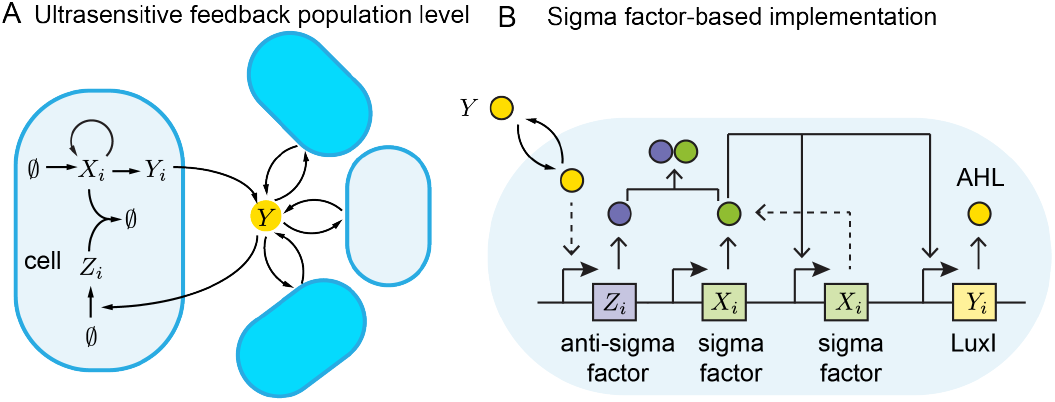
Closed-loop system with ultrasensitive feedback control. Panel A is the full circuit design with bistable switches and quorum sensing across the population. Panel B illustrates a synthetic circuit implementation using sigma and anti-sigma factors and quorum sensing molecules AHL.

We assume the AHL concentration is proportional to the enzyme LuxI. Assuming AHL reaches quasi-steady state with fast diffusion, we do not specify the intracellular and extracellular concentrations. We also assume AHL activates anti-sigma factor following a Hill-type kinetics. Then we write down the model of the closed loop of *n* cells:

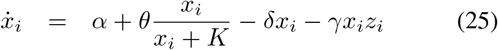

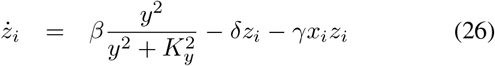

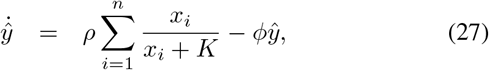

for *i*= 1: *n*. Equation (27) describes the dynamics of the total expression, which is the population-level process. According to the definition of *u* in equation (15), we can derive the steady state input-output map of the populationlevel process:

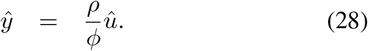

In the closed loop, the population-level controller takes *Ŷ* as the input, since *Ŷ* activates *Z_i_*’s production. We can numerically compute for the input-output map of the controller (*û* versus *ŷ*) by replacing *β* with 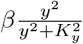 in the previous input-output map (*û* versus *β*). Then the intersection of inputoutput maps should determine the steady state equilibrium of the closed-loop system.

### A. Reference Tracking Performance

We first test if the population-level output *Ŷ* tracks different references. The references are set by the external induction, represented by parameter *α* in the model. We set three different references by increasing *α*.

We run a stochastic simulation of *n =* 100 cells in parallel and plot the time trajectory in Fig. 6A. The populationlevel output *Ŷ* (blue line) closely tracks each reference (dashed red line). Fig. 6B shows input-output maps of the population-level process and controller under corresponding reference. The process (gray line) shows a linear inputoutput map, as derived in equation (28), and the controller (orange line) exhibits an ultrasensitive input-output response. The threshold of the controller’s input-output map is moved towards the right when *α* is set with a larger value. We find that the closed-loop trajectory of *Ŷ* (blue line) indeed converges to the intersection of input-output maps of the process and the controller. The equilibrium determined by the intersection falls in the neighborhood of the threshold, which is consistent with the previous analysis of the controller.

**Fig. 6.**
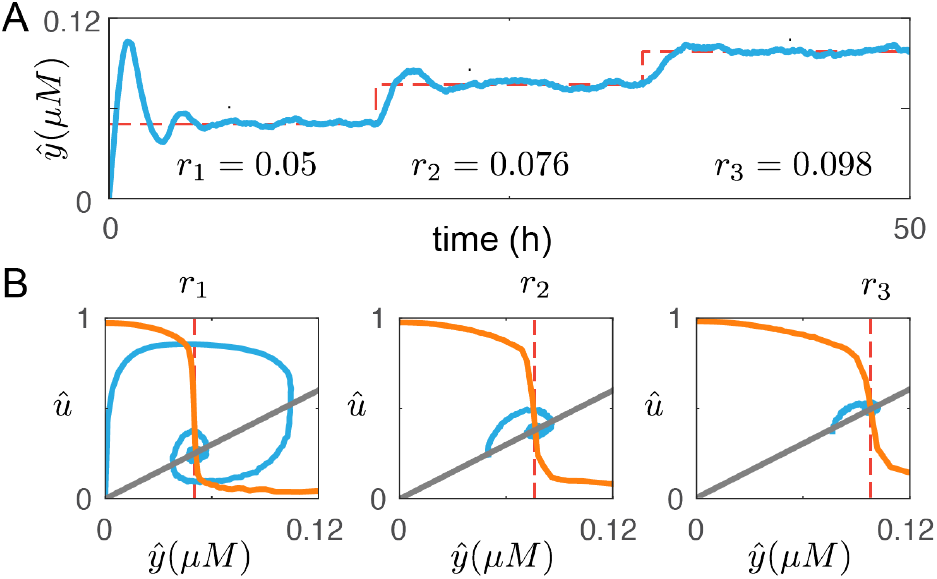
Robust tracking of references. A shows the tracking of three different references in the closed-loop system. Panel B illustrates that tracking trajectories (blue line) converge towards the equilibrium determined by the intersection of the controller’s input-output map (orange line) and the process’s (gray line) input-output map.

### B. Disturbance Rejection Performance

Next, we test if the closed-loop system can adapt to disturbances in the process dynamics via the ultrasensitive controller. We consider step disturbances that perturb the production rate *ρ* and degradation and dilution rate *φ* of *Ŷ*.

In Fig. 7A, the time trajectory shows the population-level output can adapt to disturbances with very small errors. It is more clear in Fig. 7B that the ultrasensitive controller ensures the output *Ŷ* to converge to the same concentration even with large changes in the process due to disturbances. The gray lines in the middle and right panels illustrate how the process is disturbed with a smaller ρ and a larger *φ,* compared to the left panel. Moreover, if we look at the expression state distribution for each condition in Fig. 7C, more cells switch to oN state to adapt to the disturbance, indicating the ON/OFF ratio change fulfills the ultrasensitive feedback at population-level, as demonstrated in equation (24).

**Fig. 7.**
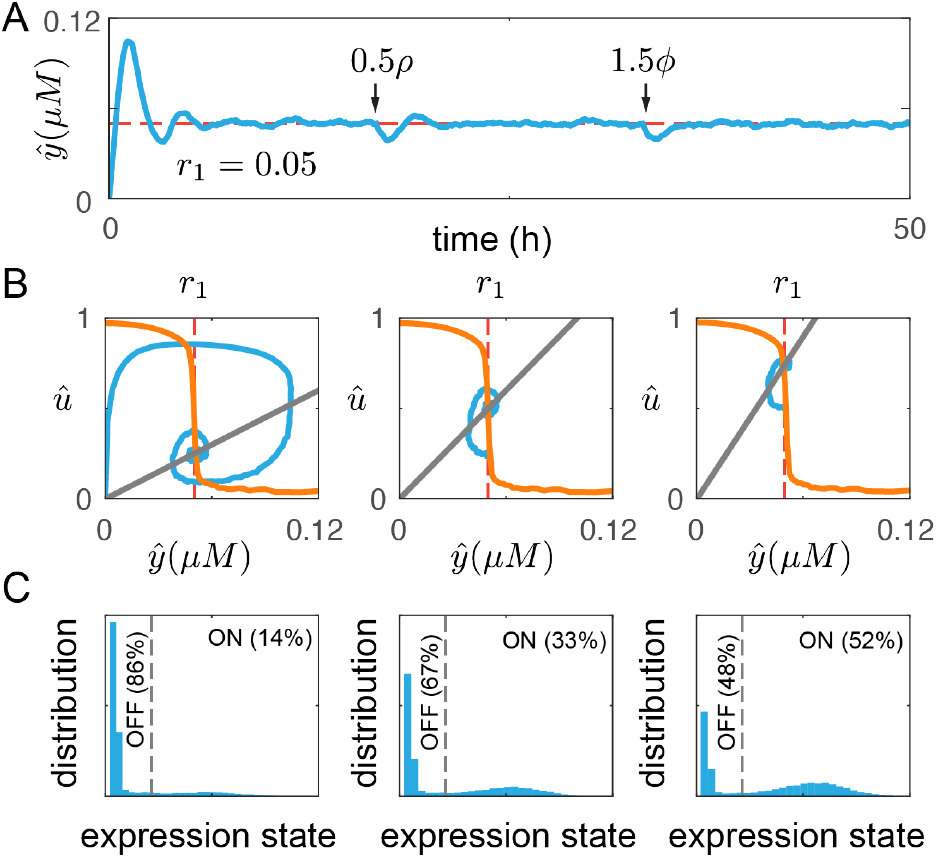
Robust adaptation to disturbances in the process dynamics. Panel A shows the adaptation of the closed-loop system when the process undergoes disturbances such as a decreased production rate *ρ* and a increased of degradation and dilution rate *φ.* Panel B shows input-output mappings of the controller and the process. All intersections are within the ultrasensitive regime, so the trajectories all converge to the same output determined by the threshold. Panel C illustrates the adaptation is achieved at population-level by changing the ratio of cells in ON and OFF states.

In summary, with the bistable switch circuit and quorum sensing system, the population-level expression robustly tracks references that are set externally and adapts to disturbances in the process dynamics. In this circuit, we consider the quorum sensing signal concentration as the target population expression. More generally, any species of interest can be controlled by having the target gene and quorum sensing signal production both activated by the controller. In that case, the quorum sensing signal can be approximated as a proportional measurement of the target expression. In addition, we can use the activator mode of the controller for more diverse implementations. Instead of having Xi activates *Y_i_* and *Ŷ* activates *Z_i_, *X*_i_* can be a repressor of *Y_i_* and *Ŷ* can be designed to activate *X*_i_ to close the negative feedback loop. In the activator mode, external induction in *Z_i_* production can set the reference, which refers to *β* in the model.

## VI. DISCUSSION

In this paper, we described how population-level ultrasensitive feedback control realizes robust and tunable target expression. The controller is based on a bistable switch resulting from a positive loop with sequestration. The quorum sensing system coordinates these switches and ensures stable and robust population-level dynamics.

In theory, we are able to predict the tunability in population-level outputs by characterizing single cell-level hysteresis. The analytical results set criteria of parameters and constraints on corresponding regulation network structures to achieve bistability. Besides the bistable switch presented in this paper, other circuits that have ultrasensitive responses such as toggle switches, phosphorylation cycles, recombinase protein switches can also be rewired with quorum sensing systems to form an ultrasensitve feedback controller at population-level.

The key design strategy of ultrasenstive controllers at population-level proposed in this paper is state switching. The required ultrasensitive response of the controller is fulfilled by sharp state switching that leads to a significant population ratio change between heterogeneous phenotypes. We demonstrate such behaviors with computations, yet more theoretical work is needed to understand what conditions guarantee an ultrasensitive response at the population-level. Here we emphasize that heterogeneity at the population-level means uniform protein overproduction is not needed in all cells: state switching results in “labor division” and reduces burden. Follow-up work will examine specifically how our strategy can improve colony survival by burden reduction, resilience to stress, and stress-related mutations.

As synthetic biological systems become increasingly complex, design principles at population-level are needed to complement single cell-level approaches. In this context, the adoption of cell-cell communication systems like quorum sensing holds great potential in achieving population control and cellular coordination, as shown by recent studies aiming at population density control [32], [33]. Cell-cell communication can also improve population-level robustness through an intercellular layer of feedback [34]. Thus, we foresee that strategies like the one described here will enable engineered microbial consortia to achieve more complex functionalities, by combining cell-cell communication and feedback architectures.

## Acknowledgements

The authors would like to thank Fangzhou Xiao, Ronghui Zhu and Nicholas A Delateur for their insightful discussions. The author XR is sponsored by the Defense Advanced Research Projects Agency (Agreement HR0011-17-2-0008). The authors EF and CCS acknowledge support by NSF/BBSRC award 2020039. The content of the information does not necessarily reflect the position or the policy of the Government, and no official endorsement should be inferred.

## Appendix

### A. Derivation for bistability conditions

We can find the steady state expression by making all equation (2)-(3) equal to zero. This results in

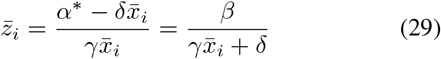

where 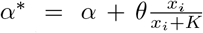. This results in a third order polynomial

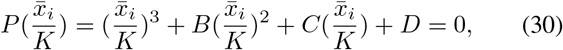

where 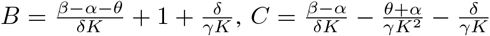, and 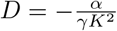.

We use the general solution of a third order polynomial, 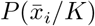, to find conditions to admit three real solutions. This leads to

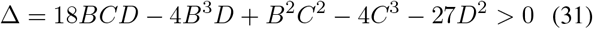

On the other hand, 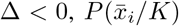 can only admit a single real solution. We consider the case when 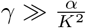, making *D* ≈ 0. Then we can simplify the condition on the number of real solutions as 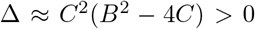. This leads to

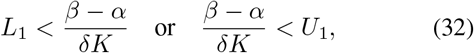

where

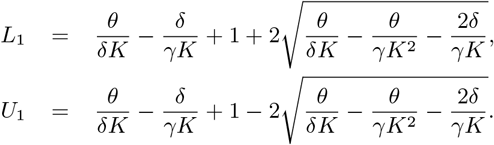

This condition does not tell us about the sign of the three real solutions. Next, we use Descartes’s rule of sign of 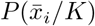 to admit three positive real solution. We need three sign changes in (1, *B,C, D).* Thus, we require *B* < 0, *C* > 0 and *D* < 0, resulting in

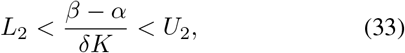

where 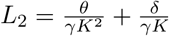, and 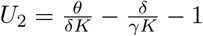.

